# Macropinocytosis and Clathrin-Dependent Endocytosis Play Pivotal Roles for the Infectious Entry of Puumala Virus

**DOI:** 10.1101/694208

**Authors:** Sandy Bauherr, Filip Larsberg, Annett Petrich, Hannah Sabeth Sperber, Victoria Klose, Walid Azab, Matthias Schade, Madlen Luckner, Chris Tina Höfer, Maik Joerg Lehmann, Peter T. Witkowski, Detlev H. Krüger, Andreas Herrmann, Roland Schwarzer

## Abstract

Viruses from the taxonomic family *Hantaviridae* are encountered as emerging pathogens causing two life-threatening human zoonoses: hemorrhagic fever with renal syndrome (HFRS) and hantavirus cardiopulmonary syndrome (HCPS) with case fatalities of up to 50%. Here we comprehensively investigated entry of the Old-World Hantavirus, Puumala virus (PUUV), into mammalian cells, showing that upon treatment with pharmacological inhibitors of macropinocytosis and clathrin-mediated endocytosis, PUUV infections are significantly reduced. We demonstrated that the inhibitors did not interfere with viral replication and that RNA interference, targeting cellular mediators of macropinocytosis, is able to decrease PUUV infection levels significantly. Moreover, we established lipophilic tracer staining of PUUV virus particles and showed co-localization of stained virions and markers of macropinocytic uptake. Cells treated with lysosomotrophic agents were shown to exhibit an increased resistance to infection, confirming previous data suggesting that a low pH-dependent step is involved in PUUV infection. Finally, we observed a significant increase in the fluid-phase uptake of cell infected with PUUV, indicative of a virus-triggered promotion of macropinocytosis.

**Author Summary:** The *Hantaviridae* family comprises a very diverse group of virus species and is considered an emerging global public health threat. Human pathogenic hantaviruses are primarily rodent-borne. Zoonosis is common with more than 150,000 annually registered cases and a case fatality index of up to 50%. Individual hantavirus species differ significantly in terms of their pathogenicity, but also their cell biology and host-pathogen interactions. In this study, we focused on the most prevalent pathogenic hantavirus in Europe, Puumala virus (PUUV), and investigated the entry and internalization of PUUV virions into mammalian cells. We showed that both, clathrin-mediated endocytosis and macropinocytosis, are cellular pathways exploited by the virus to establish productive infections and demonstrated that pharmacological inhibition of macropinocytosis or its targeted knockdown using RNA interference significantly reduced viral infections. We also found indications for an increase of macropinocytic uptake upon PUUV infections, suggesting that the virus triggers specific cellular mechanisms in order to promote its own internalization and facilitate infections.

## Introduction

Virus entry occurs through a number of successive steps that ultimately enable transferring of the pathogen’s genetic material into the host cell and hijacking of the cellular metabolism for subsequent virus replication and release.

Hantaviruses, as many related members of the order of *Bunyavirales*, have been suggested to predominantly depend on integrins(1) as cellular receptors and on endocytosis and vacuolar acidification for efficient cell entry(2). However, the specific mechanisms and processes that are involved in their internalization by host cells have only been insufficiently characterized. It was previously shown that the Old World Hantavirus, Hantaan virus (HTNV) uses clathrin-mediated endocytosis (CME)(3). Andes viruses (ANDV) on the other hand, a New World Hantavirus, was demonstrated to enter cells through clathrin-independent pathways(4). A recent study revealed that specific host kinases are required for ANDV, but also for HTNV entry, thus providing first indications for an involvement of macropinocytosis (MPC)(5). Moreover, cholesterol, which is a crucial component of many cellular processes including endocytosis, has not only been reported to be involved in entry of Crimean-Congo hemorrhagic fever virus(6), but was also demonstrated to be required for ANDV infection(7). Based on these findings it has been surmised(8) that hantaviruses, like several members of other families of bunyaviruses(9, 10), can use different pathways to finally get access to the cytoplasm of their target cells.

The canonical hantavirus cell entry mechanism entails internalization, followed by a passive trafficking within early endosomes and subsequently late endosomes. In these vesicles a pH dependent conformational change occurs that enables binding of the fusion loop of the hantavirus glycoprotein Gc to the vesicular membrane, mediating fusion between the viral and the cellular membrane and allowing release of the viral content into the cytoplasm(8). It was shown that the virus’ genetic material, together with the nucleoproteins is then transported to the site of viral replication, most likely the ER-Golgi intermediate compartment (ERGIC), via dynein-dependent trafficking along the host cell microtubules(4, 11).

In this study, we determined the cell entry mechanisms and host factors that are involved in the initial infection steps of the Puumala virus (PUUV). We utilized inhibitors of specific endocytic pathways to pinpoint productive routes of the hantavirus uptake in different mammalian cell lines by quantitative PCR. We demonstrated a cholesterol-, actin- and microtubule dependency of PUUV entry. Furthermore, we showed by confocal and correlative light and electron microscopy (CLEM) that CME and MPC are significant factors of PUUV cell entry and provided further evidence for an intracellular trafficking of fluorescently labeled viral particles along specific endocytic routes. In addition, we investigated whether cellular changes are induced by hantavirus infection that promote fluid-phase uptake by macropinocytosis using flow cytometry and confocal microscopy.

## Material and Methods

### Cell culture and PUUV infection

All adherent mammalian cell lines Vero E6 (ATCC, USA), MDCK II (ATCC, USA), Fi301 (Institute of Virology, Charité Medical School), MyGla (Institute of Virology, Charité Medical School), CHO-K1 (ATCC, USA), HeLa (ATCC, USA), A549 (ATCC, USA), Hek293T (ATCC, USA) were maintained in Dulbecco’s modified Eagle medium (DMEM) containing 10 % inactivated fetal bovine serum (FBS) and 1 % penicillin-streptomycin solution (all from PAA Laboratories GmbH, Austria).The suspension cell line K562 (ATCC, USA) and the transgenic cell line K562-aVB3 were maintained in Roswell Park Memorial Institute 1640 (RPMI 1640) containing 10 % inactivated fetal bovine serum (FBS) and 1 % penicillin-streptomycin solution (all from PAA Laboratories GmbH, Austria). K562-aVb3 cells, were recently generated by Walid Azab (Institute of Virology, Freie University Berlin) by stable transfection of a aVb3 expression plasmid and thus required antibiotic selection with 0.5 mg/ml Geneticin (Thermo Fisher, USA). If not otherwise stated, PUUV infection was performed with a multiplicity of infection (MOI) of 1 for 1 hour in medium containing 10 % FBS followed by replacement with fresh medium containing 5 % FBS.

### Virus inhibition assay

Vero E6 cells were seeded in 48-well plates and at the next day preincubated in DMEM (supplemented with 10 % inactivated FBS) with the different inhibitors (Sigma, Germany) for 1 hour: 10 mM chlorpromazine, 75 μM Ethylisopropyl amiloride (EIPA), 10 μM Rottlerin, 1 μM cytochalasin D, 20 mM ammonium chloride, 0.2 μM bafilomycin A. For methyl-β-cyclodextrin (MβCD, 10 mM) treatment, FBS free DMEM was used and replaced with FBS containing medium at 6 hours post infection (pi). 30 mM Nocodazole was applied for 2 h prior to the infection. Then, the medium was removed and the virus was added (MOI=1) diluted with the respective medium. Depending on the experimental setup, the virus mix either contained the respective compound for further inhibition of between 2 and 72 h as stated in the text or only medium without any additional inhibitor. After 24-72 h, cells were subjected to RT-qPCR analysis as described below. Virus replication was quantified by assessing the copy number of the viral S-segment per well. The inhibition efficiency was expressed as relative infection by normalizing the viral RNA copies in the inhibitor sample to the respective mock-control. Mock controls represent samples that were treated with vehicle only (DMSO or PBS depending on the solvent of the respective drug).

### Quantification of PUUV infection by RT-qPCR

Infected cells were washed with PBS, lysed with RLT-buffer and frozen at −80 °C for subsequent RNA extraction (RNA easy, Qiagen, Germany), reverse transcription and qPCR. Virus replication was quantified by assessing the copy number of the viral S-segment per well. qPCR mastermixes were prepared with 4 μl Light Cycler Enzyme Mix (Roche, Germany), 0.6 μl 10 μM primer, 0.3 μl 10 μM DNA probe, and 10.2 μl water. 4 μl of the cDNA sample together with 16 μl mastermix were combined and analyzed using a LightCycler Nano (Roche, Germany). The light cycler program was as follows: 95 °C for 10 min, followed by 40 cycles of an annealing temperature of 95 °C for 5 seconds, amplification at 60 °C for 30 seconds, and finished by denaturation at 95 °C for 15 seconds.

We used the following primers (Molbiol, Germany) and probes (Life technologies, Germany) to target the viral S-segment: PUUV F (GARRTGGACCCRGATGACGTTAA), PUUV R (CCKGGACACAYCATCTGCCAT), PUUV TMGB1 (FAM-CAACAGACAGTGTCAGCA NFQ MGB S).

### Virus labeling and purification for live cell imaging

Concentrated virus aliquots were mixed with DiI (final concentration of 1 μM, Thermo Fisher, USA), vortexed and incubated for 1.5 h at room temperature (RT). Subsequently, excess label was removed by size-exclusion chromatography using Sephadex G-25 columns (GE life sciences, USA). Labeled virus was used for visualization of internalized virus using confocal microscopy and correlative light and electron microscopy.

### Correlative light and electron microscopy

Vero E6 cells were grown and infected on microscopic dishes with finder grids (IBIDI, USA). Cells infected with fluorescently labeled virus were fixed using 4 % paraformaldehyde (PFA) in PBS and observed on a Fluoview FV-1000 confocal microscope (Olympus, Japan) to identify fluorescent spots and record their local environment. For scanning electron microscopy (SEM), cells were fixed with 2.5% (v/v) glutaraldehyde and 2% (w/v) paraformaldehyde in 100 mM cacodylate buffer (pH 7.4) for 30 min at room temperature. After fixation cells were rinsed three times for 10 min with 100 mM cacodylate buffer, and dehydrated through a graded ethanol series. After washing three times with hexamethyldisilazane (Electron Microscopy Sciences, USA) cells were coated with gold and analyzed on a LEO 1430 scanning electron microscope (Zeiss, Germany). For transmission electron microscopy (TEM), cells were fixed with 2.5% (v/v) glutaraldehyde and 2% (w/v) paraformaldehyde in 100 mM cacodylate buffer (pH 7.4) for 30 min. Cells were rinsed three times for 5 min with 100 mM cacodylate buffer postfixed for 1 h in 1% (v/v) osmiumtetroxide, rinsed three times with distilled water, en bloc stained with 0.5% (v/v) uranyl acetate, dehydrated through a graded ethanol series and finally embedded using EMBed 812 (Electron Microscopy Sciences, USA). Cells were cut perpendicular to the substrate with diamond knives on an Ultracut-R ultramicrotome (Leica, Japan) and 70–90 nm sections were collected with copper grids. Sections were counterstained with 4% (w/v) uranyl acetate followed by lead citrate. All samples were imaged on a transmission electron microscope equipped with a wide-angle CCD camera (EM 900, TRS Systems, Zeiss, Germany).

### Immunostaining, transfection and fluorescence microscopy

Vero E6 cells were seeded on glass coverslips in a 48-well plate and infected with PUUV (MOI 1) if not otherwise stated. Samples were subsequently fixed with 4 % PFA in PBS. PFA-fixed samples were permeabilized with 0.2 % TritonX-100 and stained using suitable antibodies. The viral N-protein was labeled using a primary anti-Tula virus antibody(12) and a secondary anti-rabbit FITC conjugate (Invitrogen, USA). The viral Gc protein was stained using a mouse monoclonal antibody (#H1808-60B, US Biological, US). Filamentous actin was stained with fluorescently conjugated phalloidin (Invitrogen, USA) according to the manufacturer protocol. Transfection with GFP beta-actin (addgene #27123, Addgene, USA)(13) was conducted twenty-four h prior to experiments utilizing Turbofect (Thermo Fisher, USA) according to the manufacturer’s protocol. If not otherwise stated, an inverted FluoView 1000 microscope (Olympus) was used for confocal microscopy. Differential interference contrast (DIC) and fluorescence intensity were obtained with a 60x oil immersion objective (numerical aperture 1.35) at 25 °C with a frame size of 512 × 512 pixels. Processing and analysis were carried out in imageJ (https://imagej.nih.gov/ij/).

### Spinning disk confocal microscopy

Spinning disk confocal microscopy was performed using VisiScope Scanning Disc Confocal Laser Microscope (Visitron, Germany) equipped with a 60x water objective and a EM-CCD camera.

### RNA interference

Transfection with siRNAs was performed using HiPerFect transfection reagent according to the manufacture’s reverse transfection protocol (Qiagen). First, indicated siRNAs were spotted into 48-well plates. Second, 18.5 μl of serum-free DMEM containing 1.5 μl HiPerFect was added to the pre-spotted siRNA and the mixture incubated for 20 min at RT. Afterwards, 2×10^4^ Vero E6 cells were seeded into each well, to give a final siRNA concentration of 20 nM and cells were grown under normal growth conditions for 48 h. The non-targeting siRNA AllStars (Qiagen) was used as negative control. PAK-1 targeting siRNA was purchased from Ambion (Thermo Fisher, USA).

Prior to macropinocytosis uptake experiments, transfected cells were serum-starved in DMEM supplemented with 0.1 % FCS for 3 h to promote dextran uptake. Afterwards, cells were inoculated using FITC-dextran (MW 40,000, ThermoFisher, USA) diluted in DMEM containing 0.5 % FCS to a final concentration of 4 mg/ml for 15 min at 37°C. To stop dextran uptake, cells were transferred to 4°C and washed once with cold acidic buffer (150 mM NaCl, 25 mM NaOH, pH 4) to remove any surface-bound dextran. Subsequently, cells were washed twice with cold PBS, harvested by scraping and precipitated by centrifugation (1200 × g, 5 min, 4°C). Finally, FITC intensity was determined using a FACSAria II flow cytometer (Becton Dickinson, USA). 10,000 cells were analyzed by flow cytometry.

### Flow cytometry and quantification of fluid phase uptake

To evaluate macropinocytosis upon virus infection, cells were incubated with high-titer virus solutions for 30 min at 4 °C to ensure virus binding without internalization. After removal of the virus supernatant, the cells were treated with 5 mg/ml FITC dextran (MW 40,000, ThermoFisher) for 30 min at 37 °C to target macropinosomes, washed, fixed and subjected to flow cytometry analysis. Typically, 10,000 cells were analyzed with a FACSCalibur flow cytometer (BD Biosciences, USA) and the intensity of fluorescence was analyzed using FlowJo software (Treestar, USA).

## Results

### Several mammalian cell lines are highly refractory to PUUV infections. Vero E6 cells exhibit high, MDCK-II cells modest PUUV permissivity

As a first step, we identified cell lines that are suited to study PUUV entry. An ideal model cell line would be derived from either rodent or human tissue and phenotypically resemble the natural host cell type that is pathogenically relevant in the context of hantavirus infections. As a minimal requirement, the cells should express putative hantavirus receptors and enable efficient PUUV infections. We infected a set of cell lines from different mammalian source species. Among others we tested: African Green Monkey kidney cells (Vero E6), Madin-Darby Canine Kidney (MDCK) cells; human epithelial lung (A549) cells, Henrietta Lacks (HeLa) cervical cancer-derived human cells, human embryonic lung fibroblast (Fi301) cells; Chinese hamster ovary (CHO-K1) cells; human embryonic kidney (Hek293T) cells and Mygla - a novel rodent cell line derived from the natural host of PUUV, *Myodes glareolus*.

Initially, all cell lines were infected with PUUV (MOI=1) for 120 h and analyzed by immunofluorescence staining against the viral nucleoprotein N (Figure 1). We observed a strong N protein staining in Vero E6 cells, and to a markedly lower extend in MDCK-II cells. Small numbers of infected cells were found in Fi301, Mygla, Hek293T, and A549 samples, however, N protein-positive cells were rare and typically did not show indications for productive or spreading infections. In order to verify and confirm our immunofluorescence results, we conducted another infection experiment (MOI=1) and probed virus infection by RT-qPCR at different time points post infection (p.i.). In addition to the cell lines used before, we tested another recently developed cell line that stably expresses the putative hantavirus receptor αVβ3 Integrin (k562-aVb3), as well as the respective parental cell line (K562). Of note, only the cell lines MDCK-II and Vero E6 showed significant permissiveness to PUUV infections within the time period assessed (Figure 2). Consistently, a high concentration of viral RNAs was found in Vero E6 and only a modest, significantly lower viral load was detected in MDCK-II cells. A weak virus proliferation was observed after 120 h in Mygla cells, but all other tested specimen turned out to be not significantly permissive for virus replication.

**Figure 1:**
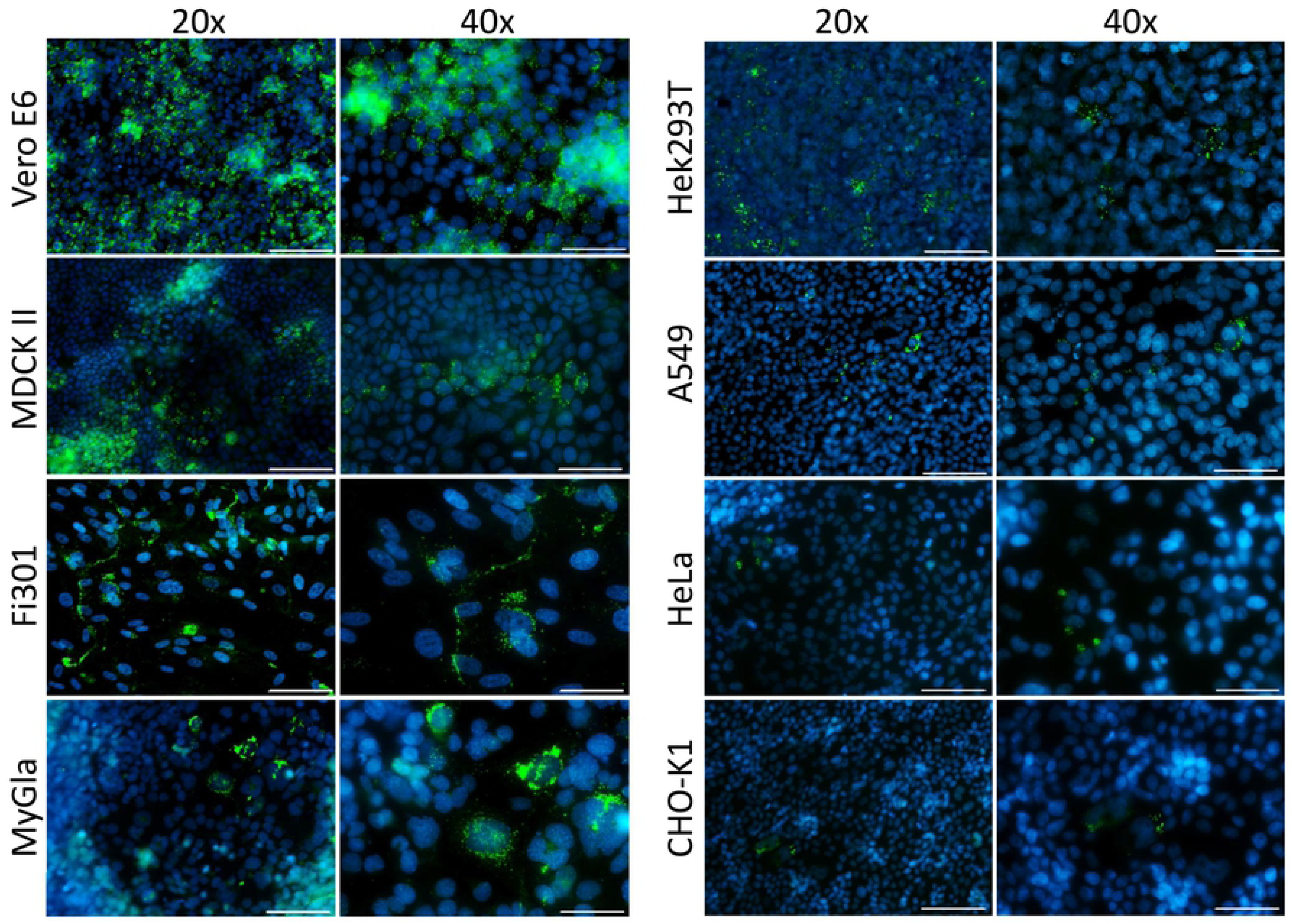
Immunofluorescence staining of PUUV N-protein in different mammalian cell lines. Cells were infected for 120 h (MOI=1) and subsequently fixed, permeabilized and stained for the PUUV nucleoprotein N (green) and DAPI as a nuclei marker (blue). Images show epifluorescence microscopy micrographs. Scale bar marks 100 μm for 20x and 50 μm for 40x magnification.

**Figure 2:**
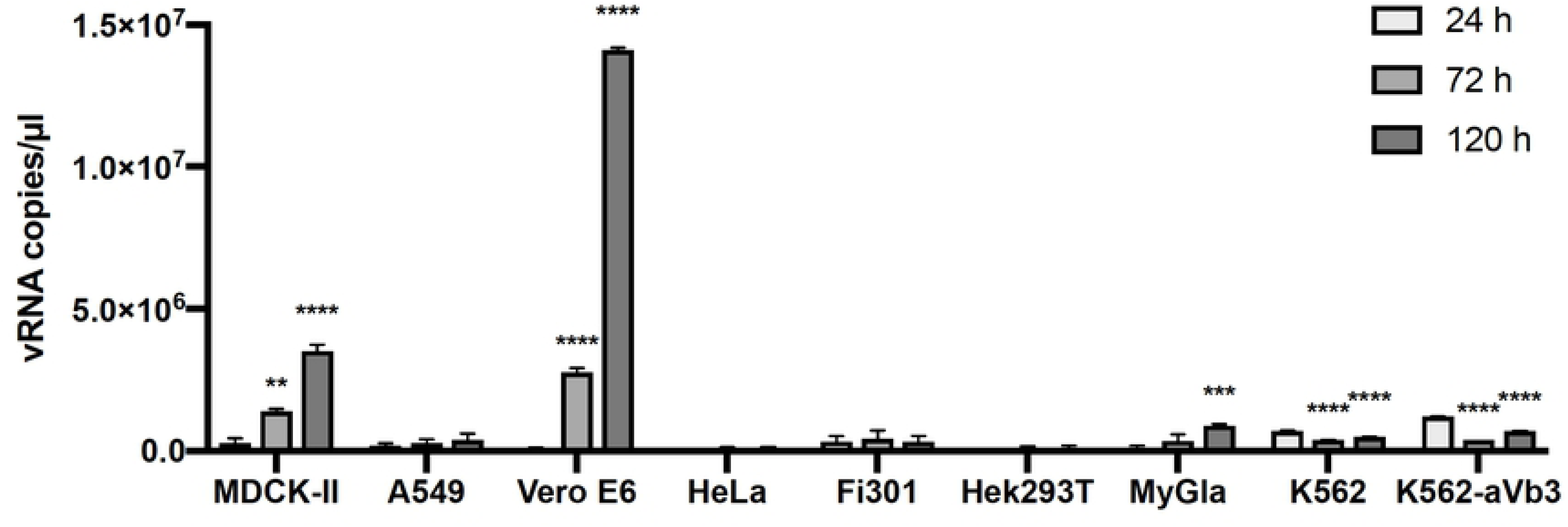
PUUV infection in different mammalian cell lines assessed by RT-qPCR. The permissivity of different mammalian cells to PUUV (MOI=1) infections was assessed by quantifying hantavirus RNA copies at different time points post infection. RT-qPCR was used to obtain the copy number of the viral S-segment. The results are based on 4 independent experiments and error bars show SEM. Significance between 24 h and 72 h or 120 h respectively, was analyzed using unpaired student’s test. **** p≦0.0001, *** p≦0.001, ** p=0.001-0,01.

Next, we stained all tested cell lines for the previously published hantavirus receptors, α5β1 integrin and αVβ3 integrin, and assessed their expression levels by fluorescence microscopy and flow cytometry (Figure S1). Of note, αVβ3 integrin expression was markedly higher in Vero E6 and MDCK-II cells as compared to all other cell lines. The alternative receptor α5β1 integrin was also expressed on non- or weakly permissive cell lines such as CHO-K1 or HeLa. This may indicate that the integrin αVβ3 is the major hantavirus receptor and predominantly responsible for productive PUUV infections. However, other cellular factors, such as antiviral, innate immunity factors, differentially expressed in specific cell lines may be involved in the permissivity as well. Consequently, we focused on Vero E6 and MDCK-II cells for all subsequent PUUV entry investigations.

### Inhibitors of clathrin-mediated endocytosis, macropinocytosis and endosomal acidification suppress PUUV infection

To unravel if PUUV utilizes different productive routes of cell entry, we used a set of nine chemical compounds known to suppress different endocytosis-related cellular mechanisms. We treated Vero E6 cells with non-toxic concentrations of the inhibitory compounds, infected them with PUUV (MOI=1) and assessed virus infection by reverse transcription quantitative PCR (RT-qPCR) at 72 h p.i. In this initial experiment, we chose a long infection time in order to ensure that detected viral RNAs (vRNA) are reflective of newly synthesized viral transcripts resulting from a successful, productive infection and subsequent virus replication. All compounds were pre-tested for inhibitory activity and cytotoxicity in Vero E6 cells and used in non-toxic, functional concentrations (Figure S2 and S3).

First, we used four compounds that affect broadly cellular internalization and trafficking mechanisms and are thus not specific for particular virus entry routes: Nocodazole, a neoplastic agent that depolarizes microtubules; Cytochalasin D, a highly active inhibitor of actin polymerization; Dynasore, a specific inhibitor of the dynamin-dependent scission of endocytic vesicles; and methyl-beta-cyclodextrin (MβCD) to disrupt plasma membrane lipid rafts by cholesterol depletion. We observed a modest (∼30 %), but significant decrease of the infection level upon treatment with either inhibitor (Figure 3A). Considering that microtubules and actin are involved in different endocytosis pathways as well as in the subsequent intracellular, endosomal trafficking(14, 15), and based on the fact that lipid rafts and cholesterol, in general, have been implicated in several endocytic pathways(16-19), these findings support and/or confirm an endocytic uptake of PUUV.

**Figure 3:**
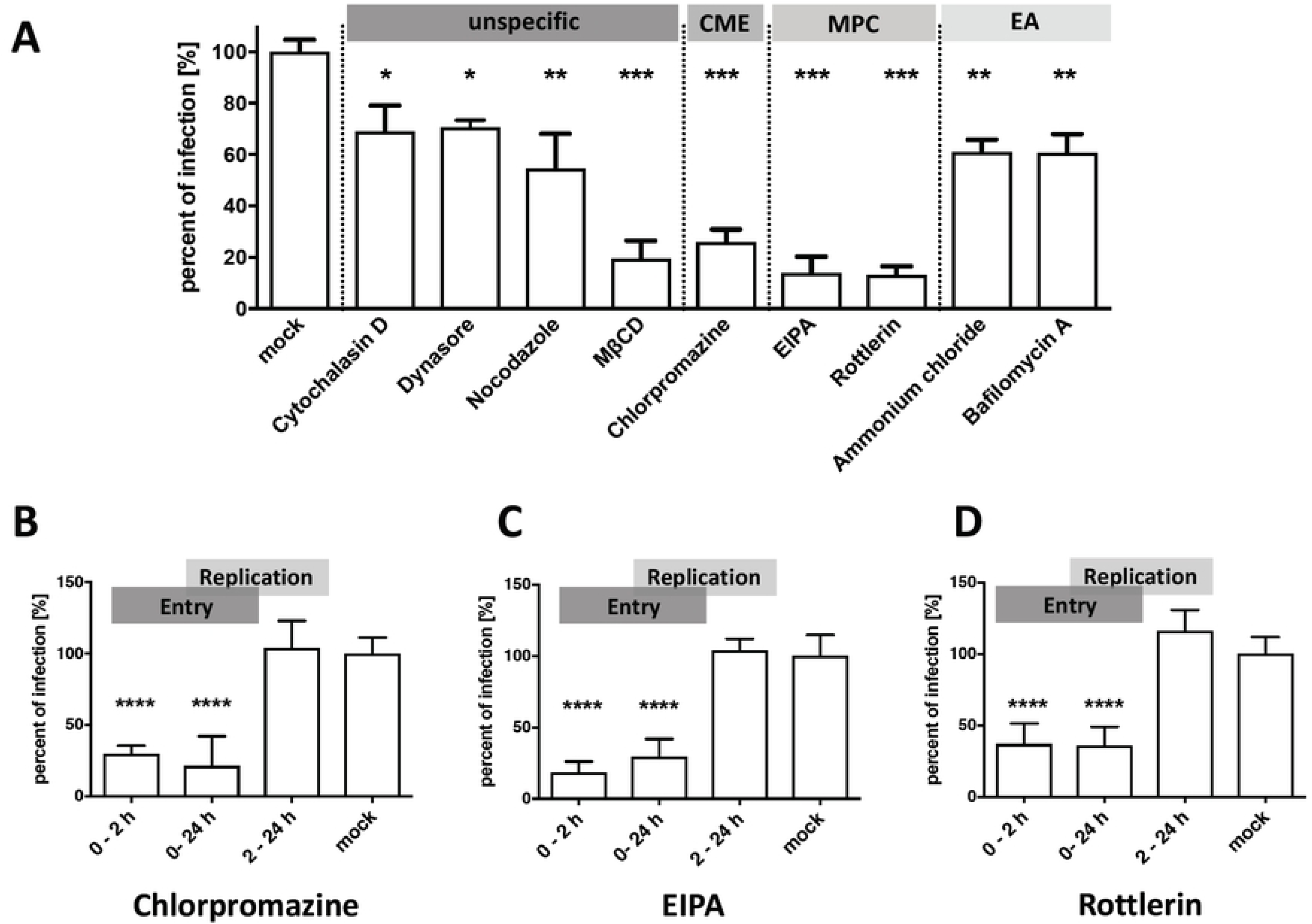
Pharmacological inhibition of virus entry into Vero E6 cells. (A) Vero E6 cells were treated with different pharmaceutical compounds to block specific virus uptake pathways and cellular endocytosis mechanisms. Cells were exposed to effective concentrations of Nocodazole (30 mM, microtubule organization), Cytochalasin D (1μM, actin polymerization), methyl-beta cyclodextrin (10 mM, MβCD, cholesterol dependent endocytosis), chlorpromazine (10 mM, initiation of clathrin mediated endocytosis), EIPA (75 μM) and Rottlerin (affecting macropinocytosis), ammonium chloride (20 mM, neutralizing acidic pH in compartments) or Bafilomycin A(0,2 μM, preventing acidification of endosomes)) and infected (MOI=1) for 72 h. CME=Clathrin mediated endocytosis, MPC=Macropinocytosis, EA=endocytic acidification. Infection levels were assessed by RT-qPCR and bars show copy numbers of the viral S-segment from inhibitor-treated samples in percent of the respective vehicle control (mock). (B-D) Transient inhibitor treatment was conducted to study the role of different compounds in PUUV entry (0-2 h p.i.) and replication (2-24 h p.i.). All bars represent at least 3 independent experiments with SEM. **** p≦0.0001, *** p≦0.001, ** p=0.001-0,01 and * p=0.01-0.05. Significance was analyzed using unpaired student’s test.

To pinpoint the biological relevance of individual cellular routes in the infection process we then used drugs targeting specific entry pathways. The drug chlorpromazine(20) has been extensively used to block clathrin-mediated endocytosis by inducing translocation of clathrin and adaptor complex 2 from the cell surface. In our experimental setup, chlorpromazine strongly reduced infection levels down to around 25%. Of note, a comparable decrease was observed with two compounds, EIPA and Rottlerin, that interfere with macropinocytosis by two different mechanisms (Figure 3A). EIPA inhibits a Na^+^/H^+^ exchanger that is involved in the formation of macropinosomes, whereas Rottlerin specifically blocks a crucial isoform of protein kinase C(21). Our finding strongly suggests that macropinocytosis is not only a bypass for PUUV infection but instead a major entry route.

Finally, Bafilomycin A, a highly specific inhibitor of a vacuolar-type H^+^-ATPase and ammonium chloride, a membrane permeable drug that buffers intracellular acidification, were used to block a putative, pH-dependent membrane fusion in cellular endosomes. Again, both compounds reduced virus infection significantly, albeit to a lower extent than EIPA and Rottlerin (Figure 3A). Taken together these experiments indicate that PUUV can enter host cells via at least two different routes. While the route via clathrin-dependent endocytosis was long known from other hantaviruses, internalization via macropinocytosis has been observed only recently for HTNV and ANDV(5).

### EIPA, chlorpromazine, and Rottlerin do not interfere with PUUV replication

To validate our previous findings, we asked whether the used chemical compounds affect viral entry, viral replication or both. (Figure 3B, C, D). Focusing on three inhibitors, we modified our previous experiments as follows: Firstly, the infection time was reduced to 24 h to prevent any influence of secondary infections (virus released from cells initially infected) on the experimental readout. Secondly, different stages of the infection process were targeted with the compounds by varying the treatment periods. Infected cells were treated with inhibitors only concomitant with the initial two hours of infection thus predominantly blocking entry, but not viral replication. Other samples were drug exposed during the whole infection time of 24 h, covering both entry and replication phase. Finally, in other samples, the treatment was started after the initial two hours of infection so that replication, but not the viral entry is affected by the compounds. Of note, a significant reduction of the infection was achieved for all three inhibitors when present in the first two hours of infection, but not if the compounds were applied after the initial entry period (Figure 3B, C, D). This finding demonstrates that the inhibitors predominantly affect the entry step and strongly support our hypothesis of a clathrin- and macropinocytosis-dominated infection of Vero E6 cells by PUUV. Of note, upon testing the CME inhibitor chlorpromazine and the MPC inhibitors Rottlerin and EIPA in MDCK-II cells, we obtained results similar to our Vero E6 data. Again, either inhibitor blocked infection efficiently when applied during the initial two hours of infection, but not if administered subsequently (Figure S4).

### Transcriptional silencing of macropinocytosis inhibits PUUV entry

Many pharmacological compounds utilized to block specific cellular pathways exert pleiotropic effects, thus affecting not only the anticipated cellular target, but also other unrelated proteins. In order to further validate our finding that inhibition of macropinocytosis suppresses PUUV entry into Vero E6 cells, we knocked-down p21-activated kinase (PAK-1), a key regulator of MPC(22) by RNA-dependent transcriptional silencing. First, we titered PAK-1 specific siRNA in Vero E6 cells, demonstrating a robust suppression of fluid-phase uptake at low nanomolar concentrations (Figure 4A). Then, cells were transfected with 20nM of PAK-1 siRNA prior to infection with PUUV, yielding significantly lower vRNA copy numbers (<80%) compared to non-targeting siRNAs. This finding confirms our pharmacological inhibition data.

**Figure 4:**
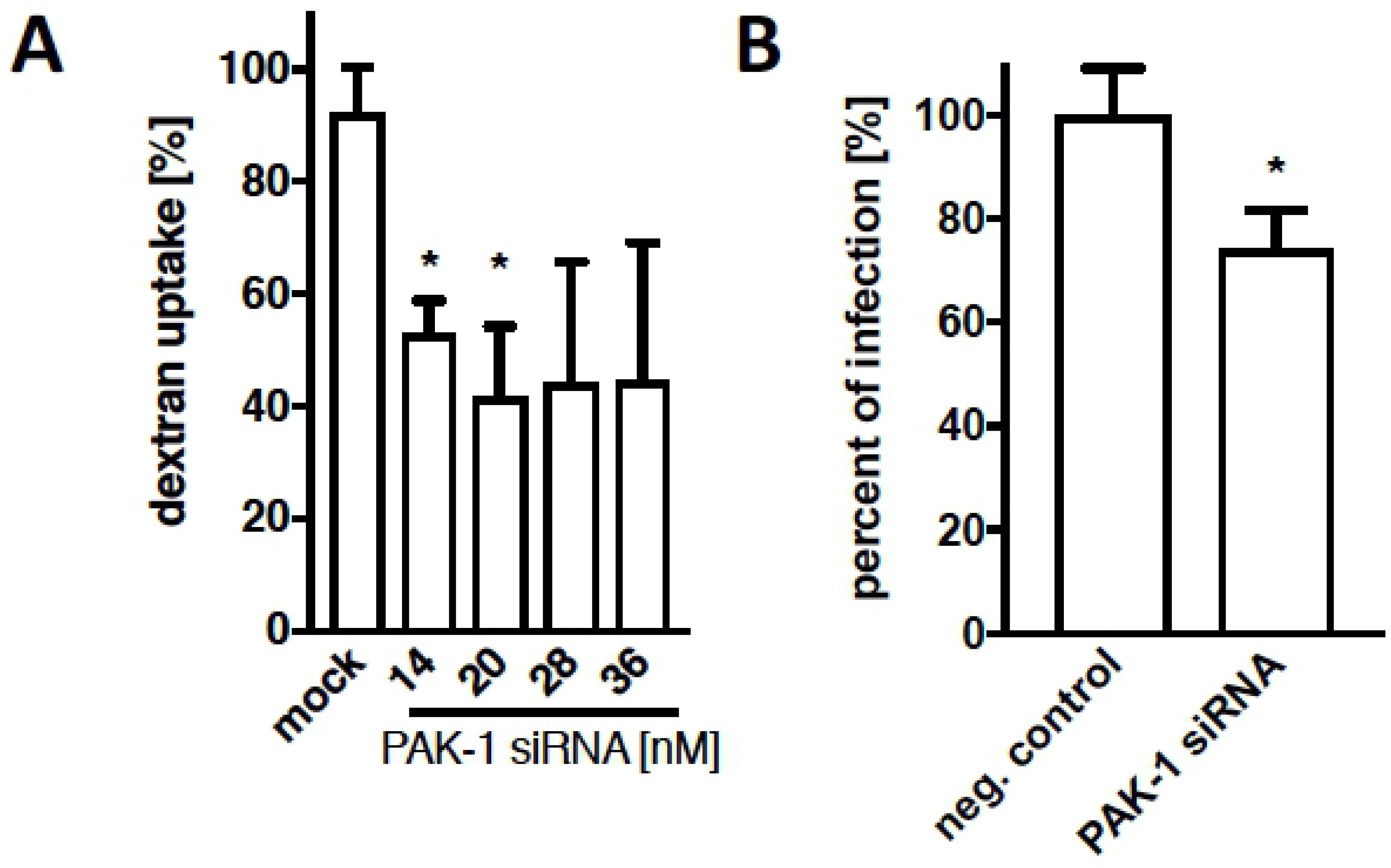
Inhibition of PUUV entry via macropinocytosis by using PAK-1 siRNA. (A) Vero E6 cells were transfected with indicated concentrations of siRNAs, targeting PAK-1 transcripts for 48 h. Subsequently, samples were incubated with FITC-dextran for 15 min and subjected to flow cytometry in order to assess fluid-phase uptake by macropinocytosis. (B) Vero E6 cells transfected with 20 nM PAK-1 siRNA for 48 h were infected with PUUV (MOI=1) for 24 h and analyzed for hantavirus infection levels. Viral RNA copy numbers were assessed by RT-qPCR and bars show results from inhibitor treated samples (in %) of the respective non-targeting siRNA. All bars represent 3 independent experiments with SEM. Significance was analyzed using unpaired student’s test, * p<0.05. Mock and negative control show samples transfected with non-targeting siRNA.

### Lipophilic tracer staining of PUUV enables live virus detection by fluorescence microscopy

Hitherto, macropinocytosis has not been implicated in PUUV entry. In order to provide independent evidence for a PUUV uptake into host cells by macropinocytosis, we therefore established live-cell imaging of PUUV entry as a complementary approach. To enable tracing of internalized virions we used a virus membrane-staining approach that has been previously used for various enveloped viruses(23).

Virus preparations were treated with the lipophilic dye DiI and subsequently purified using size-exclusion-chromatography. Stained virus particles were then subjected to live cell, confocal microscopy and observed over a period of 30 to 60 min. In cells, previously transfected with a fluorescently labeled microtubule binding protein (Tau-YFP) we were able to show directed particle trafficking along cytoskeleton components (Fig. 5A). We analyzed 5 particles trajectories and found an average particle velocity of around 1,000 nm/sec (see supplementary video), which is in agreement with a kinesin/dynein-mediated transport along cellular microtubule filaments (24).

**Figure 5:**
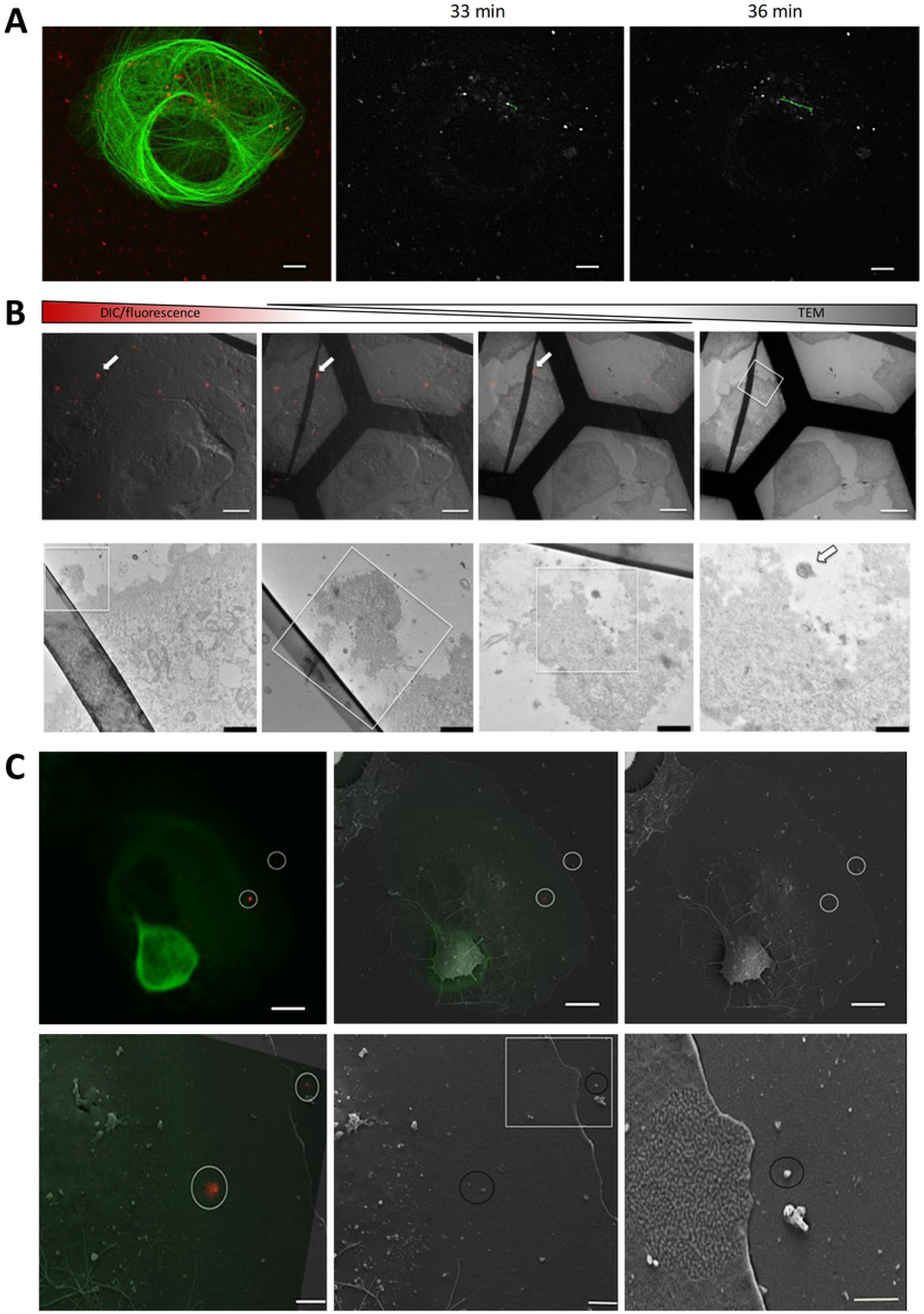
PUUV particles labeled with lipophilic tracers enable live fluorescence microscopy and correlative light and electron microscopy. (A) Vero E6 cells transfected with Tau-YFP were infected with DiI labeled virus particles and subjected to live confocal microscopy. Images were acquired every 2.9 sec and virus particles were tracked using ImageJ. Micrographs show overlays of Tau-YFP expression (green) and virus particles (red) in the left image, as well as virus particles (in grey) at different time points post infection in the central and right image. A representative trajectory of a single virus particle is shown in in the right image (green). (B and C) Correlative light and electron microscopy (CLEM) was used to reveal ultrastructures of stained particles. First, samples were analyzed by confocal microscopy, recording specific regions of interest in gridded microscopy slides. Subsequently, the same samples were analyzed by electron microscopy to unravel the ultrastructure of previously observed particles. (B) The upper panel shows a gradual overlay of differential interference contrast/fluorescence microscopy and Transmission electron microscopy images. Scale bar 5μm. The lower panel displays magnifications of the boxed regions. Scale bar 2, 0.6, 0.4, 0.25μm, respectively from left to right. (C) The upper panel shows a representative fluorescence microscopy image on the left, the corresponding region in a scanning electron microscopy image on the right and an overlay of both in the middle. Fluorescent particles are highlighted with white circles. Scale bar 5μm. The lower panel shows magnifications of the regions containing fluorescent particles. Scale bars: 2μm, 2μm and 1μm from left to right.

To validate our labeling procedure, different controls were performed. First, cells infected with labeled-hantavirus were subjected to Correlative Light and Electron Microscopy in order to show that intracellular, fluorescent spots were representative of virus particles and not artifacts, such as fluorophore aggregates or stained cell-debris. In fact, fluorescent spots were recovered in regions showing spherical particles in raster electron microscopy (Figure 5C). Moreover, transmission electron microscopy (TEM) revealed electron-dense virions (Figure 5B) that coincide spatially with fluorescent particles in infected Vero E6 cells. Of note, the electron microscopy images show a typical viral nanostructure with a diameter of around 100 nm in regions that were correlated with their respective counterparts in the confocal images based on surrounding cellular structures and topographies. We also tested whether labeled viruses were still able to productively infect their host cells or otherwise, whether internalized virions were rendered non-infective by the labeling procedure. To this end, we infected Vero E6 cells with labeled viruses and assessed virus infection using RT-qPCR at 48 h post-infection. We found robust and high levels of infection, demonstrating that the staining and washing procedure did not abrogate the infectivity of the viral particles (Figure S5).

### PUUV particles co-localize with macropinocytic cargo

Next, we asked if hantavirus particles can be found in macropinosomes upon cell entry. To this aim, Vero E6 cells were incubated concomitantly with virus and high-molecular weight dextran, both fluorescently-labeled. Fluorescent dextran is an established marker of macropinocytosis(25) and spatial correlation of virus and dextran indicates a joint uptake by macropinosomes. We found colocalization between FITC-dextran and the lipophilic tracer DiI as early as 10 min after initiation of the experiment (not shown), but a more robust intracellular signal of both fluorescent marker was obtained at 30 min post infection (Figure 6). In this experiment, the samples were fixed, additionally stained for the PUUV glycoprotein Gc, the viral N protein or both, and imaged by spinning disc confocal microscopy. A high degree of colocalization was found for Gc and the virus membrane marker DiI (Figure 6A), thus confirming that lipophilic tracer staining enables efficient and specific detection of PUUV particles. Moreover, all viral markers showed distinct co-localization with dextran-containing compartments, representative of a macropinocytic uptake of PUUV particles (Figure 6A, B). Of note, we also observed internalized virions, which did not co-localize with FITC-dextran containing vesicles. Dextran colocalization negative virus signals may indicate particles that have already left the macropinocytic pathway or reflect virus entry through alternative endocytic pathways, i.e. clathrin mediated endocytosis.

**Figure 6:**
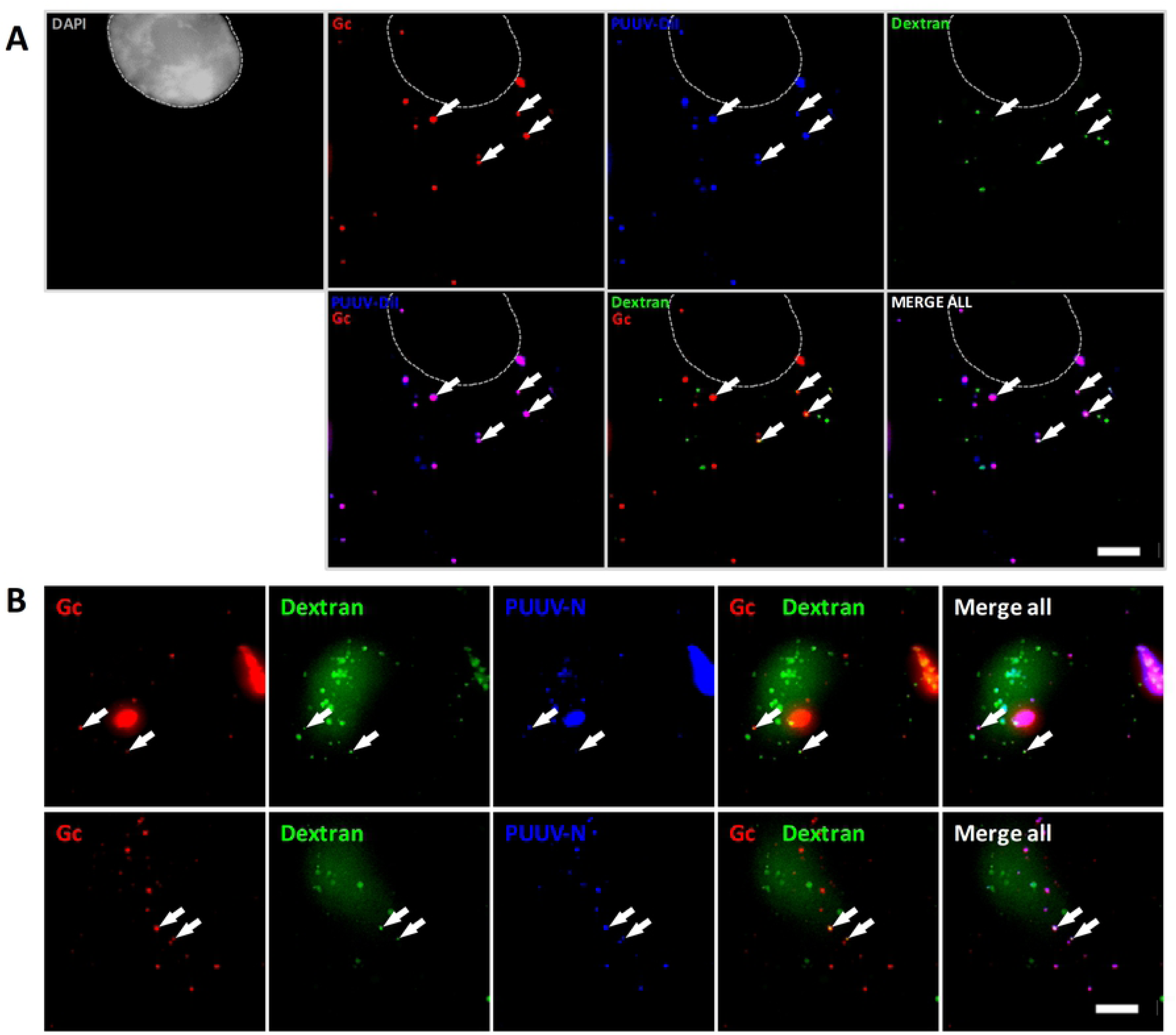
Colocalization of macropinocytic cargo and PUUV. (A and B) DiI-labeled viral particles were used to infect Vero E6 cells, concomitantly exposed to FITC-dextran in order to specifically label macropinosomes. At 30 min post infection, cells were fixed and stained with DAPI and antibodies against PUUV Gc. (B) Cells fixed after 30 min dextran-uptake, stained with Gc and N antibodies. Arrows indicate viral particles that colocalize with dextran-containing compartments. Samples were imaged using spinning disk microscopy. Scale bar 5 μm.

### Hantavirus infection promotes the formation of actin foci and activates macropinocytosis

Hantaviruses have been previously reported to utilize cytoskeleton components in order to promote intracellular particle trafficking (4). In addition, we have demonstrated that labeled PUUV virions travel along host microtubules (Figure 5A) and found indications for a kinesin/dynein-mediated transport. Here, we sought to investigate whether actin contributes to the intracellular PUUV lifecycle. Actin is a key factor for the formation of membrane ruffles and macropinosomes, but also critically involved in other cellular endocytosis mechanisms (26). In order to study cellular actin in the context of PUUV infections, we stained infected cells for the viral N-protein and filamentous actin (F-actin) using Rhodamine-Phalloidin at 72 h post infection.

Interestingly, we often observed an increased accumulation of F-Actin in regions and particularly in cells with high concentrations of the viral N-Protein (Figure 7A, solid arrows) as compared to non-infected cells (Figure 7A, dashed arrows, Figure S6, Figure S7). We surmised that this region harbors the microtubule organizing center (MTOC) and therefore performed an additional Tubulin immunofluorescence staining. In fact, viral enrichment was often found in the vicinity of perinuclear microtubule aggregates (Figure 7B, Figure S8). Both observations support previous reports about the importance of the cytoskeleton in hantavirus infections (4, 27). Of note, other virus families have been previously reported to stimulate endocytosis pathways and in particular macropinocytosis in order to facilitate their uptake and productive entry into host cell (28, 29). Based on this notion and our previous results, we hypothesized that PUUV infections may increase the uptake of high molecular-weight dextran. To test this hypothesis, we infected cells with high virus titer (MOI=10) for 30 min at 4 °C to enable binding without internalization and membrane fusion. After removal of the virions, FITC-dextran was used to quantify fluid-phase uptake via flow cytometry. We found a slight, but significant increase of FITC-dextran uptake in virus samples compared to mock-infected negative controls in each of 5 independent experiments (Figure 7B), indicating that macropinocytic endocytosis is enhanced upon PUUV receptor binding.

**Figure 7:**
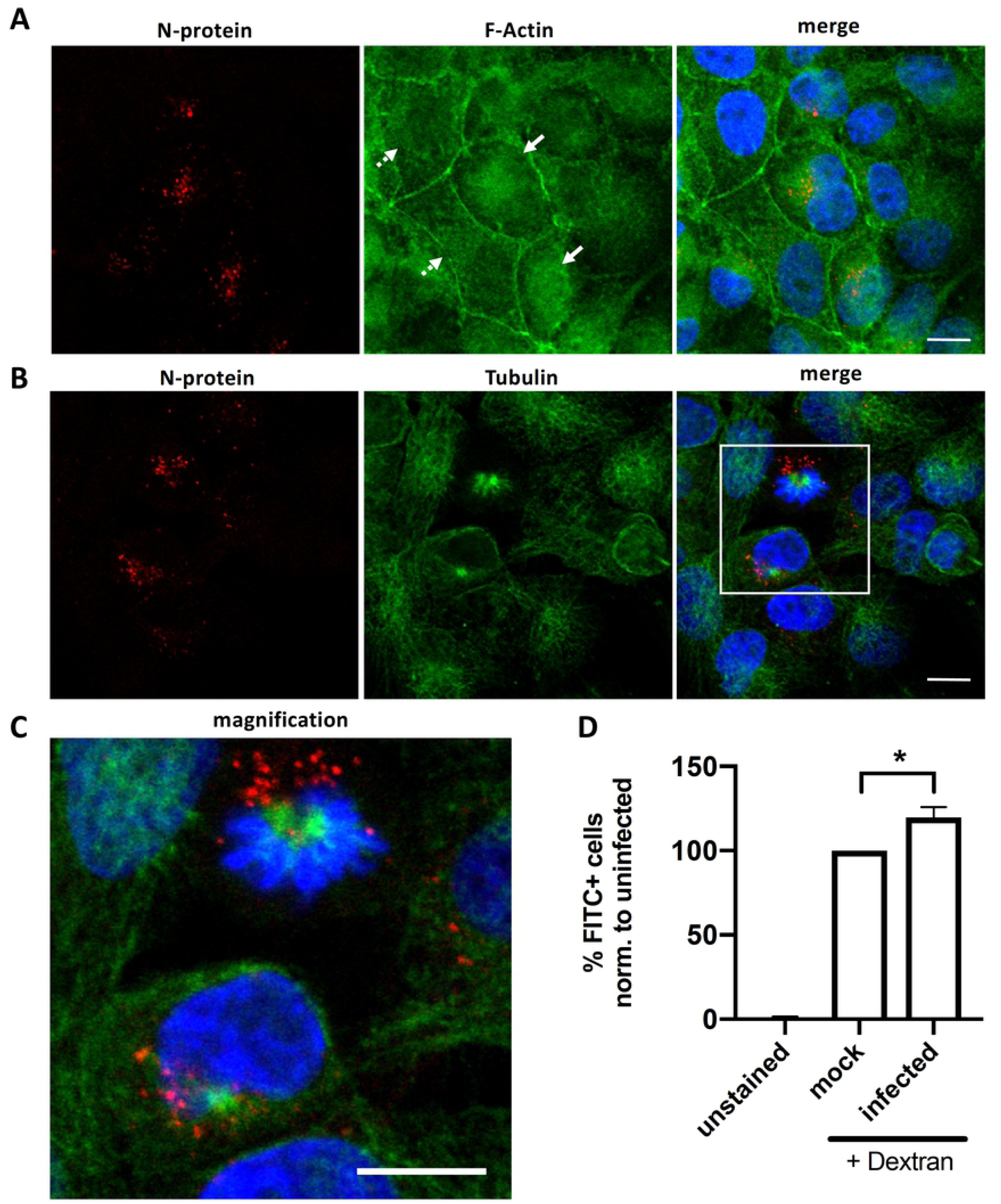
Stimulation of fluid-phase uptake by PUUV. (A and B) Vero E6 cells infected with PUUV for 72 h, and stained for (A) filamentous actin using Rhodamine-Phalloidin or (B) microtubules using ***α***-tubulin and N-protein, the latter of which were assessed by immunofluorescence. White arrows indicated infected, dashed arrows highlight uninfected cells. (C) Magnification of the boxed are in B. Scale bars: 10 μm. (D) Vero E6 cells, infected with hantavirus (MOI=10) for 30 min at 4 **°**C were subsequently investigated using flow cytometry for FITC-dextran internalization as a reporter of fluid-phase uptake. Bars represent 5 independent experiments with SEM. Significance was analyzed using unpaire student’s test,* p≦0.05.

## Discussion

Natural hosts of hantaviruses are small rodents, shrews, moles and bats. Secondary infections have not been reported in other mammals, but are frequently found in humans (30, 31). In order to identify a cell line model that is as physiologic and yet sufficiently permissive for hantavirus infections, we initially screened immortalized cell lines from different species that are commonly available in most research facilities. Upon infection with PUUV we observed that only two of them Vero E6 and MDCK-II displayed increasing and high levels of viral RNA and proteins (Figure 1 and 2) within the first 5 days of infection. The bank vole derived cell line MyGla also supported productive infections albeit delayed as compared to MDCK-II and Vero E6 cells. Interestingly, we consistently observed a significant decrease of the PUUV RNA levels in K562 cells, which may indicate rapid and autocrine antiviral response to hantavirus infections in this cell line. Of note, surface expression levels of the putative hantavirus receptors αVβ3 integrin, but not α5β1 integrin seem to correlate with the permissivity of the different cell lines (Figure S1).

Aim of this study was to investigate the cellular mechanisms and pathways that permit PUUV entry and enable successful infection of mammalian cells. In recent years, hantaviruses have been reported to enter their host cells via different pathways of receptor-mediated endocytosis. For instance, the Old World hantavirus HTNV was shown to be internalized by clathrin-mediated endocytosis (CME)(3), whereas other members of the hantavirus family were demonstrated to be internalized through CME-independent route(4). Recently, Torriani et al.(5) utilized a pseudo virion reporter system to investigate entry of both Old and New world hantaviruses. Noteworthy, the authors provided first evidence for a contribution of macropinocytosis to the entry of both Old and New world hantaviruses. Here we utilized replication-competent PUUV particles to unequivocally study the early steps of the viral life cycle.

Upon treatment of Vero E6 cells with inhibitors specific either for clathrin-mediated endocytosis (chlorpromazin) or for macropinocytosis (EIPA, Rottlerin) we found a significant reduction in infection levels, suggesting that PUUV is able to utilize both clathrin-mediated uptake and macropinocytosis for virus entry and infection (Figure 3). Moreover, we found that cholesterol depletion with MßCD significantly reduced PUUV infection. Cholesterol is a major structural element of mammalian membranes and has been shown to be involved in numerous endocytic mechanisms(18, 32, 33). Consistent with our finding, cholesterol has been previously shown to be required for the infection of Andes virus(ANDV)(7) as well as other virus species(34, 35).

Hantavirus receptor binding and membrane fusion are exclusively mediated by the glycoprotein complex (GPC), consisting of the two subunits Gn and Gc. The latter, Gc has been proposed to constitute a class II fusion protein(36), thus requiring acidification-induced conformational changes in order to elicit membrane fusion. Our finding of a significant reduction of hantavirus infections with inhibitors of intracellular acidification is coherent with this hypothesis (Figure 3).

To the best of our knowledge, an involvement of macropinocytosis in cell entry and infection of PUUV has not been reported before. Torriani et. al.(5) were the first to demonstrate that pharmacological inhibition of cellular factors, believed to be strictly required for macropinocytic internalization is sufficient to significantly suppress hantavirus uptake. This study however was focused only on HTNV and ANDV, members of the Old and New world orthohantavirus genus, respectively, and did not include other hantavirus species, such as PUUV. In addition, assays solely based on pharmacologic inhibition of cellular proteins and mechanisms are often plagued by a limited reliability of the results due to the pleiotropic nature of the used drugs. Therefore, to further validate our inhibitor based results, we sought to provide additional independent evidence for an involvement of macropinocytosis in PUUV infections. To this aim, we first employed RNA interference to transcriptionally silence PAK1, a crucial component of the cellular macropinocytosis machinery, which confirmed our finding of significant reduction of PUUV infection (Figure 4). Then, we established lipophilic tracer staining of PUUV particles to assess virus uptake by fluorescence microscopy. This method has been extensively used to study entry and membrane fusion of other viruses, such as i.e. Influenza(37), but has not yet been stringently tested and utilized in hantavirus studies. We found that labeled particles can be visualized by live-fluorescence microscopy for extended periods of time (Figure 5) and validated their ultrastructure by CLEM (Figure 5). With this system in place, we tested whether PUUV enters cells through macropinosomes, which were visualized using FITC-Dextran. Indeed, we found co-localization of FITC-Dextran and stained viral particles as well as viral proteins, detected by immunofluorescence, thus again confirming macropinocytosis as a biologically relevant cellular entry route of PUUV (Figure 6).

Cellular actin is key mediator of macropinocytic uptake and several cytoskeleton components have been reported to be utilized by different hantaviruses species(4). In agreement with that, we also observed intracellular actin accumulation in infected, but not in uninfected Vero E6 cells. Noteworthy, several virus species that utilize macropinocytosis for cell entry and infection have been shown to actively trigger and promote this endocytic mechanism(28, 29). To test whether PUUV induces macropinocytosis, we first infected Vero E6 cells and subsequently tested them for FITC-dextran uptake by flow cytometry. We found a mild, but significant elevation of the FITC-dextran after only 30 min of incubation with PUUV virus particles. In the light of the high MOI we have used here, further studies are warranted to support this notion. However, an almost identical observation was previous made for Ebola virus like particles(28), which may suggest a common virus entry mechanism that is shared by *Filoviridae* and *Hantaviridae*.

In this work, we performed a comprehensive study of productive PUUV entry pathways and provided strong indications for a contribution of macropinocytosis during PUUV infection. Noteworthy, host Integrins, which are widely believed to be the predominant receptors for hantaviruses(1) have been previously reported to traffic through macropinocytosis(38). In addition, integrin targeting antibodies were show to be internalized by macropinocytosis(39), again suggesting a direct mechanistic connection between these receptors and macropinocytic endocytosis. Future studies will have to further investigate the role and significance of different cellular entry pathways for hantavirus entry into endothelial cells, the major PUUV target cell type.

## Supporting Information Legends

### Supplementary Material

Includes SI Figures S1 to S8.

**Supplementary Video S1:** Representative VeroE6 cell labeled with Tau-YFP (video S1) and infected with DiI labelled PUUV particles (video S2). Images were acquired with a Fluoview FV-1000 confocal microscope (Olympus, Japan) using a UPLSAPO 60X Water objective (NA:1.20). Images were recorded every 6.58 sec.

**Supplementary Video S2**: Representative VeroE6 cell labeled with Tau-YFP (video S1) and infected with DiI labelled PUUV particles (video S2). Images were acquired with a Fluoview FV-1000 confocal microscope (Olympus, Japan) using a UPLSAPO 60X Water objective (NA:1.20). Images were recorded every 6.58 sec.

**Supplementary Video S3**: Overlay of video S1 and S2.

